# BAHCC1 promotes gene expression in neuronal cells by antagonizing SIN3A-HDAC1

**DOI:** 10.1101/2025.01.24.634719

**Authors:** Alan Monziani, Rotem Ben-Tov Perry, Hadas Hezroni, Igor Ulitsky

**Affiliations:** Department of Immunology and Regenerative Biology and Department of Molecular Neuroscience, Weizmann Institute of Science, Rehovot 76100, Israel

## Abstract

Chromatin modifications play a key role in regulating gene expression during development and adult physiology. Histone acetylation, particularly H3K27ac, is associated with increased activity of gene regulatory elements such as enhancers and promoters. However, the regulation of the machinery that write, read, and erase this modification remains poorly understood. In particular, the SIN3A-HDAC1 complex possesses histone deacetylase activity, yet it commonly resides at active regulatory regions. Here, we study BAHCC1, a large chromatin-associated protein essential for viability, and recently reported to play a largely repressive role. We show that in neuronal lineage cells, BAHCC1 is mainly associated with regulatory elements marked with H3K27ac. BAHCC1 interacts and co-occupies shared genomic regions with the SIN3A scaffold protein, and its perturbations lead to altered acetylation and expression of proximal genes in a neuronal cell line and primary cortical neurons. The regulated genes are enriched for those functioning in neurogenesis and cell migration, and primary cortical neurons with reduced *Bahcc1* expression display impaired neurite outgrowth. We thus propose a model in which BAHCC1 antagonizes SIN3A histone deacetylation and positively regulates the expression of genes that are important for growth and migration-related processes in the neuronal lineage.

## Introduction

Regulation of gene expression plays a central role in establishing cell type identity during embryonic development and the response of cells to external stimuli throughout their lifetime. Gene expression is regulated by combinations of regulatory elements found at promoters and enhancers, whose accessibility and activity are modulated by chromatin conformation and histone marks. Various proteins and protein complexes write, read, and erase the histone marks (1). While the activity of some of these complexes, such as the repressive Polycomb PRC1 and PRC2 complexes, has been extensively studied and is reasonably well understood, others remain largely obscure (2, 3). SIN3A and the related SIN3B proteins are at the heart of the Sin3 multisubunit complexes. The structure of the SIN3A complex has been resolved and it contains 13 core proteins with SIN3A acting as the main protein scaffold (4). Loss of function of the core members of Sin3 results in early embryonic lethality (5–8), suggesting a crucial role during early development, yet its biological function remains largely unclear. The Sin3 complex contains HDAC1, a histone deacetylase associated with gene repression. Sin3 was traditionally thought to have repressive roles (9, 10), as recruitment of SIN3A is sufficient for inducing repression at target promoters (11, 12). However, HDAC1 is also part of the NuRD and CoREST complexes, and other HDACs also deacetylate histones, so the relative contribution of SIN3A is unclear. Chromatin profiling studies located Sin3a complexes at active promoters in mouse embryonic stem cells (13) and other systems (14, 15), casting doubt on whether it regulates gene expression negatively or positively.

BAHCC1 is a poorly studied chromatin-associated protein. BAHCC1 is a ∼280 kD protein conserved throughout vertebrates and characterized by a BAH domain found at its C-terminus and a pair of Tudor domains. Its expression is up-regulated in acute lymphoblastic leukemia (ALL) of T-cell or B-cell lineages (T-ALL or B-ALL) and in melanoma, and its depletion affects the proliferation of leukemic and melanoma cells (16, 17). BAHCC1 is an ortholog of the D. melanogaster *winged eye* (wge), and a paralog of TNRC18, both of which were implicated in different gene expression regulation steps (18–20). A recent study has shown that the BAH domain of BAHCC1 binds the H3K27me3 repressive mark and suggested that BAHCC1 acts as a ‘reader’ of H3K27me3 marks and represses gene expression in blood cells (16), though most of the biochemical characterization focused on just the C-terminal BAH domain, which accounts for just ∼5% of the full protein. In contrast, BAHCC1 was predominantly associated with active H3K27ac-demarcated regions and positively regulated gene expression in melanoma cells (17).

We recently studied *Bahcc1* in mouse embryonic stem cells and early neuronal progenitors in a study focused on a neighboring gene, a long noncoding RNA called *Reno1* (21). We found that KO of *Bahcc1* or *Reno1* inhibited neuronal differentiation of mESCs and triggered cell death, consistent with a defect in neurogenesis upon knockdown (KD) of BAHCC1 in another study (22). We also found that KO of BAHCC1 led to a substantial reduction of chromatin accessibility at regions marked by H3K4me3 and the SIN3A-FAM60A complex in mouse embryonic stem cells (21). Other studies have shown that BAHCC1 co-immunoprecipitates with members of the SIN3A complex, including HDAC1, SAP30, FAM60A, RBBP4, RBBP7, and ING2 (13, 16, 23–26). Additional data point to the importance of BAHCC1 in the neuronal lineage - *Bahcc1* mRNA is preferentially expressed in the developing brain over other tissues (21), and *Bahcc1*^−/−^ mice exhibiting perinatal lethality, with one mouse reported to show defective motor neuron development (27). The requirement of BAHCC1 for embryonic development and postnatal survival was also observed in another mouse model carrying a specific mutation in the BAH domain (16). In this study, we focused on the gene regulatory roles of BAHCC1 in neuronal lineage cells.

## Results

### Bahcc1 binds active regulatory regions in a neuronal cell line

To characterize the roles of BAHCC1 in cells of a neuronal lineage, we used the mouse Neuro2a (N2a) neuroblastoma cell line, which is readily transfectable, thus permitting transient manipulations and large-scale biochemical and genomic assays. To profile BAHCC1 genomic occupancy, we used a BAHCC1 polyclonal antibody we developed (**Fig. 1A**) for a CUT&RUN (28) assay. We first tested the specificity of this antibody by Western blot upon *Bahcc1* knockdown (KD) and overexpression (OE), and by immunofluorescence (IF) after transfecting a plasmid encoding an HA-tagged full-length *Bahcc1* coding sequence (CDS). These experiments show that the BAHCC1 signal, as reported by the antibody, responds to changes in *Bahcc1* levels (**Fig. 1A**, left), and that its staining overlaps that of HA in transfected cells (**Fig. 1A**, right). In parallel, we profiled the occupancy of an HA-tagged full-length BAHCC1 protein expressed from a transfected plasmid, allowing Dox-inducible BAHCC1 expression. Both experiments revealed concordant binding profiles (ρ = 0.48, P < 10^−15^, **Fig. 1B**), suggesting that *Bahcc1* overexpression and tagging do not strongly affect its genome-wide occupancy. Therefore, we merged all conditions/libraries and plotted the coverage over metagenes and other epigenetic features. BAHCC1 was enriched immediately upstream to the TSS of active genes (**Fig. 1C**), in agreement with its profile in melanoma cells (17). BAHCC1 CUT&RUN read coverage was slightly more prominent among promoters of more highly expressed genes (**Fig. 1C**) and largely absent around those not expressed in N2a cells, suggesting a role in regulating gene expression. We also profiled the canonical chromatin marks H3K27ac, H3K4me3, and H3K27me3 in untransfected N2a cells and found that BAHCC1 peaks were enriched with both H3K4me3 and H3K27ac (**Fig. 1D-E**), displaying a higher relative signal for the latter. In contrast, BAHCC1 peaks were largely devoid of H3K27me3, in contrast to a recent report suggesting BAHCC1 is a reader of this chromatin mark (16). Because of the association of BAHCC1 with SIN3A complex members in other cell lines (13, 16, 23–26), we profiled SIN3A occupancy in N2a cells by CUT&RUN. We found that SIN3A and BAHCC1 share a similar binding profile (**Fig. 1D-E**). Both BAHCC1 and SIN3A were more strongly associated with H3K27ac-marked regions, with BAHCC1 displaying a preference for intergenic regions (**Fig. 1F**, top) and SIN3A for promoter-associated ones (**Fig. 1F**, bottom). When considering the genome-wide CUT&RUN read coverage, BAHCC1 showed the strongest correlation with H3K27ac and SIN3A, a moderate correlation with H3K4me3, and minimal correlation with H3K27me3 (**Fig. 1G**), consistent with our previous observations from the metagene profiles. Overall, BAHCC1 chromatin occupancy suggests that it is associated with regions marked with acetylated histones and the SIN3A complex, and this association increases with increasing expression levels of associated genes.

**Figure 1.**
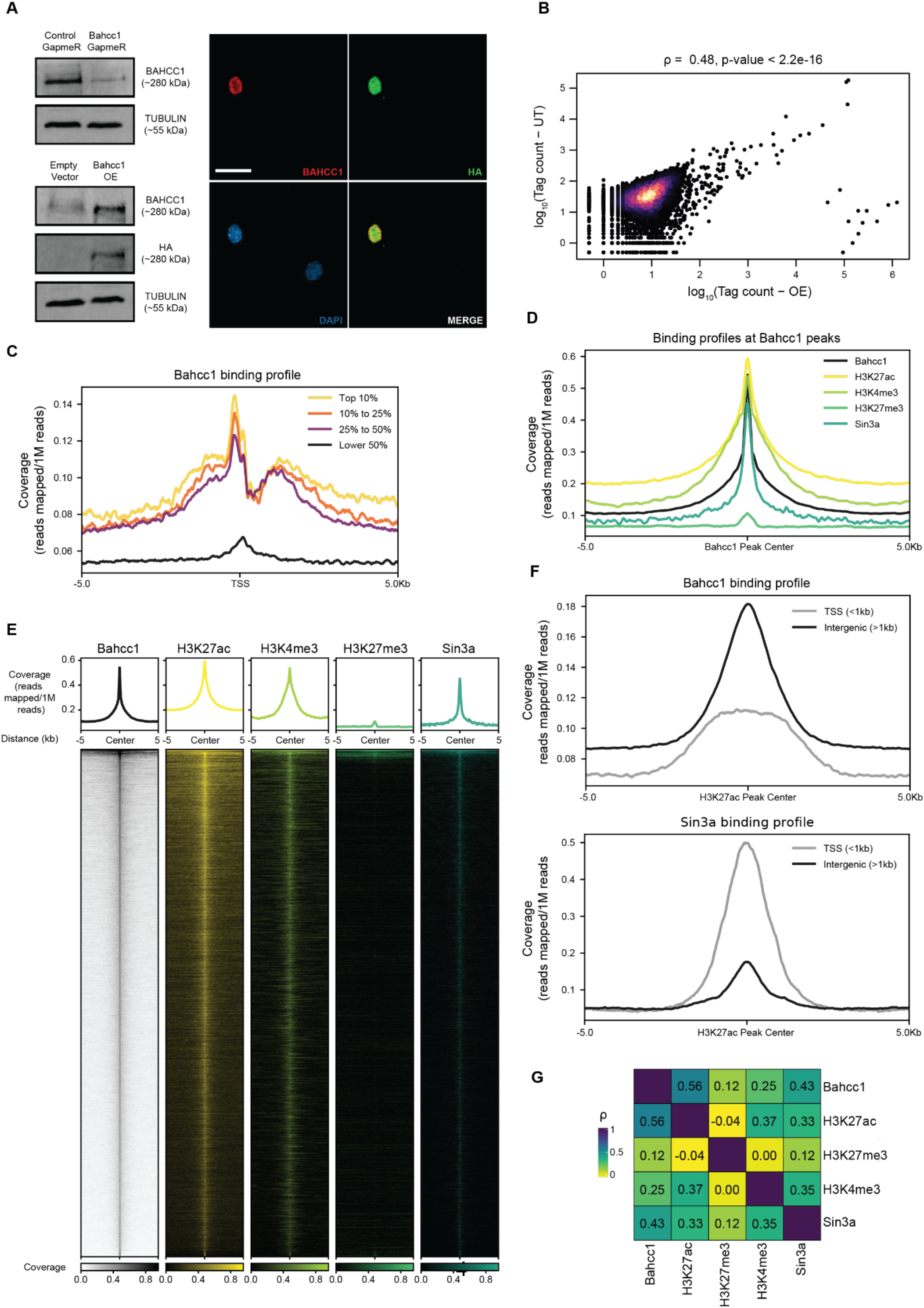
BAHCC1 chromatin occupancy in N2a cells. **(A)** Left: Western blots using the polyclonal BAHCC1 antibody developed in this study upon *Bahcc1* KD and OE; Right: IF showing co-localization of the BAHCC1 antibody signal with the HA-tag (scale bar = 20 µm, 100X magnification). **(B)** Correspondence between BAHCC1 CUT&RUN signals in untransfected (UT) and *Bahcc1*-overexpressing N2a cells. Each point is a binding peak detected by combining all the data together. Color indicates local point density. **(C)** Metagene profile around TSS of the normalized BAHCC1 CUT&RUN signal in N2a cells, stratified by gene expression levels. **(D)** Metagene profiles of normalized CUT&RUN signals of H3K4me3, H3K27ac, and SIN3A, centered over BAHCC1 peaks. **(E)** Heatmap of read coverage around the BAHCC1 peaks, ranked by BAHCC1 binding and centered around the peak. Coverage and heatmaps of BAHCC1, H3K27ac, H3K4me3, H3K27me3, and SIN3A are shown. **(F)** Metagene profiles of normalized CUT&RUN signals of BAHCC1 (top) and SIN3A (bottom), centered over H3K27ac TSS-associated (<1kb) or intergenic (>1kb) peaks. **(G)** Heatmap for the correlations of the CUT&RUN coverage between the examined proteins and histone modifications. Colors and numbers represent the Spearman’s correlation coefficients.

### BAHCC1 levels regulate gene expression in neuronal cells

To study the effect of BAHCC1 dosage, we perturbed BAHCC1 expression by different methods and quantified changes in gene expression by polyA(+) RNA-seq. Specifically, we knocked down *Bahcc1* in N2a cells using GapmeRs (29) – antisense oligonucleotides that trigger RNA degradation via nuclear RNAse H activity (**Fig. 1A** and **2A**), as we found this approach to be more effective than siRNAs for depleting *Bahcc1* mRNA in N2a cells (**Fig. S1A and S1B**). Furthermore, we knocked down *Bahcc1* in two different cellular states – undifferentiated N2a cells and cells differentiated into a more mature phenotype by a retinoic acid (RA) treatment(30) for 72 hrs (**Fig. S2A**). We have also overexpressed *Bahcc1* in undifferentiated cells by transient transfection of a plasmid encoding an HA-tagged *Bahcc1* CDS (**Fig. 1A** and **2B**). We analyzed changes in gene expression on these conditions using DeSeq2 (31), considering a gene to be differentially expressed if its adjusted P < 0.05 and it changed by at least 33% (|log_2_FC| > .41). As expected, RA-mediated differentiation caused substantial transcriptomic changes (**Fig. S2B**), with an increase in the expression of genes associated with neuronal maturation processes (**Fig. S2C**), and a decrease in those associated with cell adhesion, migration, and proliferation (**Fig. S2D**). Among the upregulated genes, several were related to the biogenesis, transport and release of dopamine, a feature previously observed in N2a cells (32).

**Figure 2.**
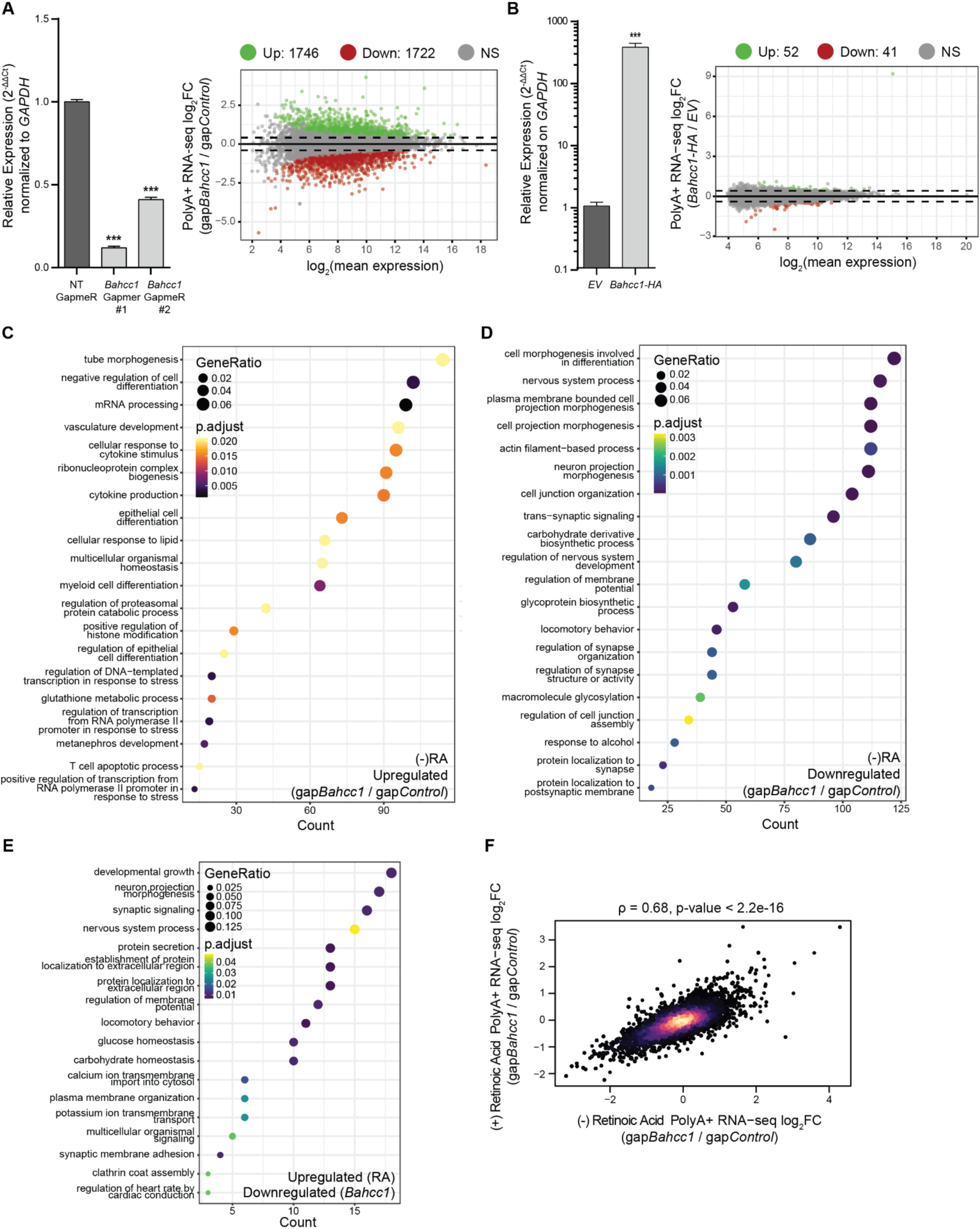
Genome-wide transcriptomic changes upon BAHCC1 perturbations. **(A)** RT-qPCR upon *Bahcc1* depletion with GapmeRs to assess KD efficiency (left) and an MA plot showing the changes in genome-wide gene expression (right). **(B)** RT-qPCR upon transient transfection of a plasmid encoding *Bahcc1* CDS with a C-terminal HA-tag to assess OE efficiency (left), and MA plot showing the changes in genome-wide gene expression (right). **(C)** GO enrichment analysis of the upregulated genes following *Bahcc1* KD with GapmeRs, in undifferentiated cells. **(D)** Same as (C), but for the downregulated genes. **(E)** GO enrichment analysis of genes normally upregulated by RA treatment, but instead repressed by *Bahcc1* depletion with GapmeRs during differentiation. **(F)** Correspondence between changes in gene expression after *Bahcc1* KD with GapmeRs in (-)RA and (+)RA N2a cells. Color indicates local point density. All experiments were performed in n = 3 biological replicates, with the error bars in the barplots representing the standard deviation, P > .05 = ns; < 0.05 = *, < .01 = **; < .001 = *** (two-tailed Student’s t-test). Genes were considered to be differentially expressed when adj. P < 0.05 and |log_2_FC| > .41 (corresponding to a change of 33%).

We then focused on genes that were dysregulated after perturbing *Bahcc1* levels. Genes upregulated following *Bahcc1* KD in either (+)RA or (-)RA cells were enriched for processes related to stress response and immune activation (**Fig. 2C** and **S2E**), whereas downregulated genes were strongly linked to neuronal development, maintenance, and functions (**Fig. 2D** and **S2F**). Genes that were significantly upregulated after retinoic acid treatment and downregulated after *Bahcc1* KD in (+)RA cells (i.e., genes that fail to be properly activated by the differentiation process) are enriched in several processes related to neuronal morphogenesis, synapsis development, and signaling (**Fig. 2E**), hinting that BAHCC1 activity could be related to neuronal differentiation. However, we observed no substantial morphological changes in the differentiating cells, and gene expression changes following *Bahcc1* KD before and after differentiation were broadly similar (**Fig. 2F**), suggesting that BAHCC1 regulates a shared set of genes regardless of the cellular state. Thus, in subsequent analysis, we focused on changes in gene expression in undifferentiated cells. Overexpression of HA-tagged *Bahcc1* had modest effects on gene expression (**Fig. 2B**). The upregulated genes were enriched for immune activation and cellular proliferation (**Fig. S2G**), whereas the downregulated one, were involved in general RNA processing (**Fig. S2H**). These results indicate that perturbing BAHCC1 levels triggers substantial changes in N2a cells that are consistent between undifferentiated and differentiated cells, nevertheless affecting genes important for neuronal development, maintenance, and function. Notably, since *Bahcc1* OE has more modest effects compared to its KD, endogenous levels of BAHCC1 in N2a cells appear to suffice to exert its functions. The enrichment of cell locomotion-associated genes among the genes downregulated following *Bahcc1* KD led us to examine cell migration and invasion in *Bahcc1* KD cells. *Bahcc1* KD resulted in a marked reduction in the ability of N2a cells to bridge the gap in a wound healing assay (**Fig. S3A-D**), as well as to migrate in a nutrient gradient as assessed in a transwell assay (**Fig. S3E**). These effects of *Bahcc1* KD are consistent with those observed in melanoma cells, where *Bahcc1* KD led to reduced cell proliferation and invasion (17). Taken together, these results indicate that BAHCC1 activity is required to establish a proper gene expression program in N2a cells, and its dysregulation causes decreased expression of lineage-commitment genes and reduced cell motility.

### BAHCC1 positively regulates gene expression by regulating H3K27ac levels at promoter-distal sites

To evaluate the effects of BAHCC1 on chromatin, we used CUT&RUN in biological triplicates to profile H3K27ac and H3K4me3 in *Bahcc1* KD and OE cells. We associated each of the 46,129 H3K27ac peaks we identified in N2a cells (**Fig. 3A**) with the nearest gene and used DeSeq2 to compute the effect of *Bahcc1* perturbations on each peak. Since changes in gene expression were overall modest, we focused first on the difference between *Bahcc1* OE and KD (comparing each to their respective controls) and compared the changes in gene expression to changes in H3K27ac and H3K4me3. Changes in gene expression were significantly correlated with changes in H3K27ac at the associated peaks (R=0.13, P<10^−15^), whereas a substantially weaker correlation was observed for H3K4me3 (R=– 0.04, P=7.2×10^−7^). The association with H3K27ac was weaker yet still significant separately for the KD and the OE of BAHCC1 (KD: R=0.08 and P<10^−15^; OE: R=0.04 and P=8.9×10^−9^).

**Figure 3.**
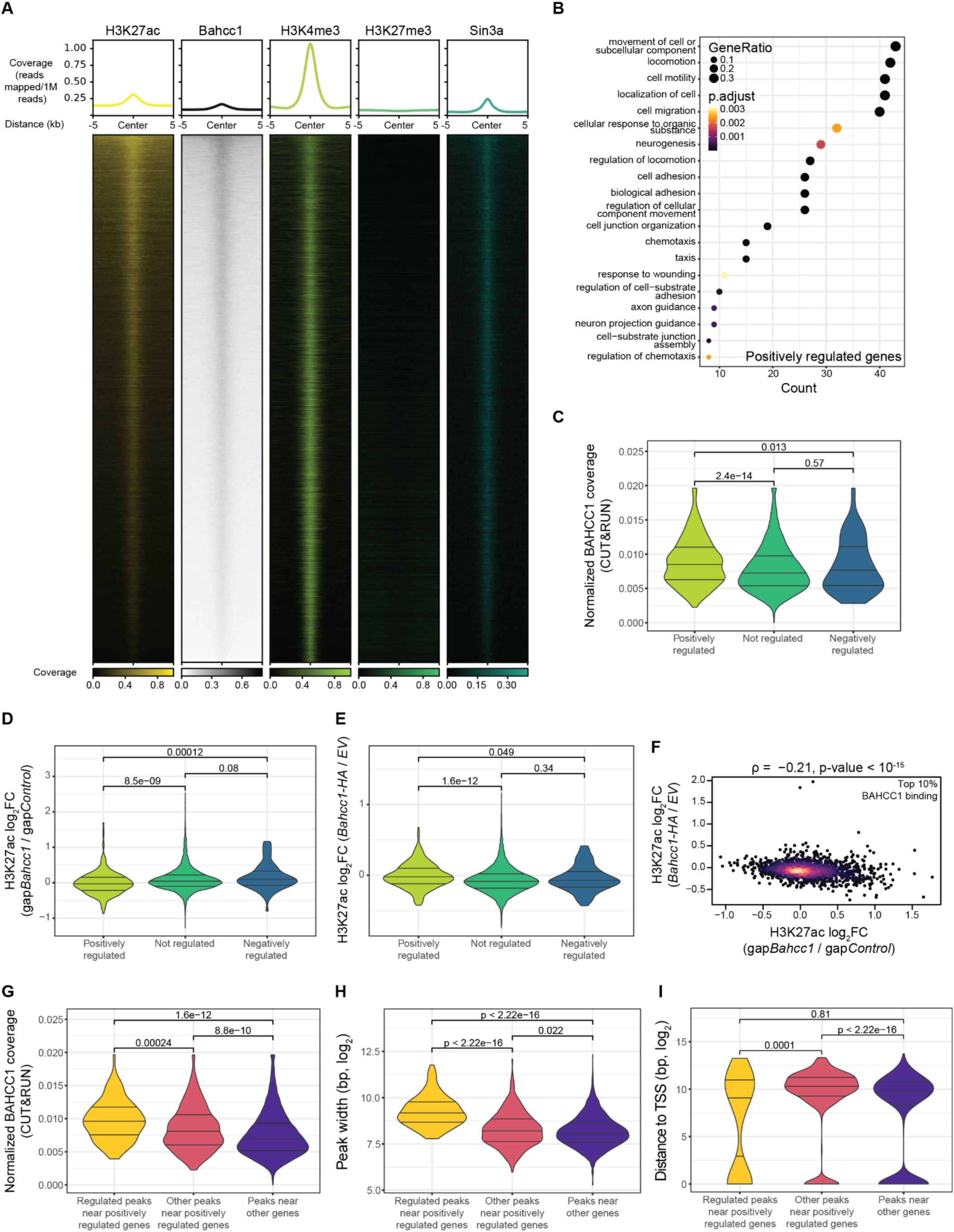
BAHCC1 positively regulates gene expression. **(A)** Heatmap of all high-confidence H3K27ac peaks, ranked by BAHCC1 coverage. Coverage of BAHCC1, H3K4me3, H3K27me3 and SIN3A, as well as their metagene profiles are shown. **(B)** GO enrichment of the positively regulated genes. **(C)** Normalized BAHCC1 CUT&RUN signal over H3K27ac peaks near positively, negatively, and unregulated genes. **(D)** Changes in H3K27ac at peaks near the BAHCC1-regulated genes in *Bahcc1* KD cells. **(E)** Same as in (D), but after *Bahcc1* overexpression. **(F)** Correspondence between changes in H3K27ac after *Bahcc1* KD and OE for the peaks with the top 10% average coverage of BAHCC1 CUT&RUN reads. Color indicates local point density. **(G)** Normalized BAHCC1 CUT&RUN signal over the H3K27ac regulated peaks associated with the positively regulated genes. **(H)** Width (in log scale) of the regulated H3K27ac peaks. **(I)** Distance from the nearest TSS of the regulated H3K27ac peaks. In all violin plots, the three lines represent the median, first and third quartiles, and significance of the different comparisons was calculated by using a Mann-Whitney test.

We then focused specifically on genes that were differentially regulated by the BAHCC1 dosage and defined 613 ‘positively regulated’ genes as those that were significantly (P<0.05) increased by *Bahcc1* OE, and significantly decreased by its KD, and had a change of at least 25% following either OE or KD. Conversely, we defined 134 negatively regulated genes as those that decreased following the OE and increased following KD, with the same criteria. Positively regulated genes were strongly enriched in biological processes related to neuron development, projection guidance, adhesion, and synaptic signaling (**Fig. 3B**). In contrast, negatively regulated ones were not enriched in any biological process. These two gene sets were compared to 10,230 ‘not regulated’ genes whose transcripts changed by less than 25%, with P>0.05 in both OE and KD. BAHCC1 occupied peaks associated with positively regulated genes more than peaks associated with the negatively regulated ones or ones that were not regulated (**Fig. 3C**). The peaks associated with the positively regulated genes had reduced H3K27ac following *Bahcc1* KD (**Fig. 3D**), whereas *Bahcc1* OE increased H3K27ac in these regions (**Fig. 3E**). Peaks associated with negatively regulated genes were instead not significantly affected (**Fig. 3D-E**). BAHCC1 KD and OE led to significantly anti-correlated changes in H3K27ac at the peaks with the 10% strongest BAHCC1 binding (R=–0.21, P<10^−15^, **Fig. 3F**), whereas in the peaks in the bottom 50% of BAHCC1 binding the anti-correlation was much weaker (R=–0.03, P=0.005, **Fig. S4A**), pointing to a direct involvement of BAHCC1 in regulation of H3K27ac levels. H3K4me3 levels were affected only by Bahcc1 KD and exclusively in those peaks associated with the positively regulated genes (**Fig. S4B-C**). No substantial anti-correlation was observed when considering H3K4me3 at BAHCC1-bound peaks (R=–0.01 P=0.36), again primarily associating BAHCC1 with H3K27ac.

We then considered the 97 H3K27ac peaks near the positively regulated genes that increased by Bahcc1 OE (P<0.05) or decreased by its KD (P<0.05). We compared them to 399 peaks that did not change by 25% in either perturbation and 7,343 peaks near not-regulated genes (P>0.05) in both conditions. The peaks in the first group exhibited higher levels of both BAHCC1 and SIN3A binding (**Fig. 3G** and **Fig. S4D**), were broader (**Fig. 3H**), and slightly closer to the transcription start sites compared to the other peaks associated with the positively regulated genes (**Fig. 3I**). Interestingly, positively regulated genes had lower SIN3A occupancy in their promoters compared to the other categories (**Fig. S4E**), suggesting that when BAHCC1 is present, SIN3A does not bind or binds with lower efficiency. Notably, SIN3A occupancy is higher at promoters of highly transcribed genes, despite being considered a co-repressor given its involvement in SIN3-HDAC1 (**Fig. S4F**). We conclude that BAHCC1 primarily promotes gene expression, and changes in its levels are associated with changes in the H3K27ac marks in regions bound by BAHCC1.

### BAHCC1 associates with SIN3A in neuronal cells

Following the reported associations between BAHCC1 and SIN3A in other cell types (13, 16, 23–26), we wanted to further confirm this by native co-immunoprecipitation (co-IP) upon the transfection of the full-length *Bahcc1* OE plasmid in HEK293T cells. Both HA and BAHCC1 staining revealed that we successfully pulled down BAHCC1, and staining with a SIN3A antibody validated the predicted interaction (**Fig. 4A**, left). Moreover, co-IP using the SIN3A antibody successfully retrieved BAHCC1 (**Fig. 4A**, right), again confirming the association between BAHCC1 and SIN3A. To get a better overview of the BAHCC1 interactome, we performed liquid chromatography followed by mass spectrometry (LC-MS) of the eluates from the co-IP using an anti-HA antibody, after transfecting the cells with either the full-length *Bahcc1-HA* or empty vector. Strikingly, all the significantly enriched (> 2 unique peptides, fold-enrichment > 2, P<0.05) proteins were part of the SIN3A-HDAC complex (**Fig. 4B**). These include structural and/or organizer proteins such as SIN3A, SAP130 and SAP30, as well as SUDS3, BRMS1 and BRMS1L which were previously found to enhance the interaction with HDAC1 and the deacetylase activity of the complex (8, 33–35). HDAC1 and RBBP7 were also found to be significantly enriched, but they did not meet the fold change criteria. Notably, SIN3B was not recovered as associated with BAHCC1. We further tested the colocalization of BAHCC1 with SIN3A and different histone marks (H3K27ac, H3K4me3 and H3K27me3) by immunofluorescence. BAHCC1 was found exclusively in the nuclei of N2a cells, where it was co-localized with H3K4me3, H3K27ac, and SIN3A (**Fig. 4C**). In contrast, we found again no evidence for co-localization of BAHCC1 with H3K27me3 (**Fig. 4C**).

**Figure 4.**
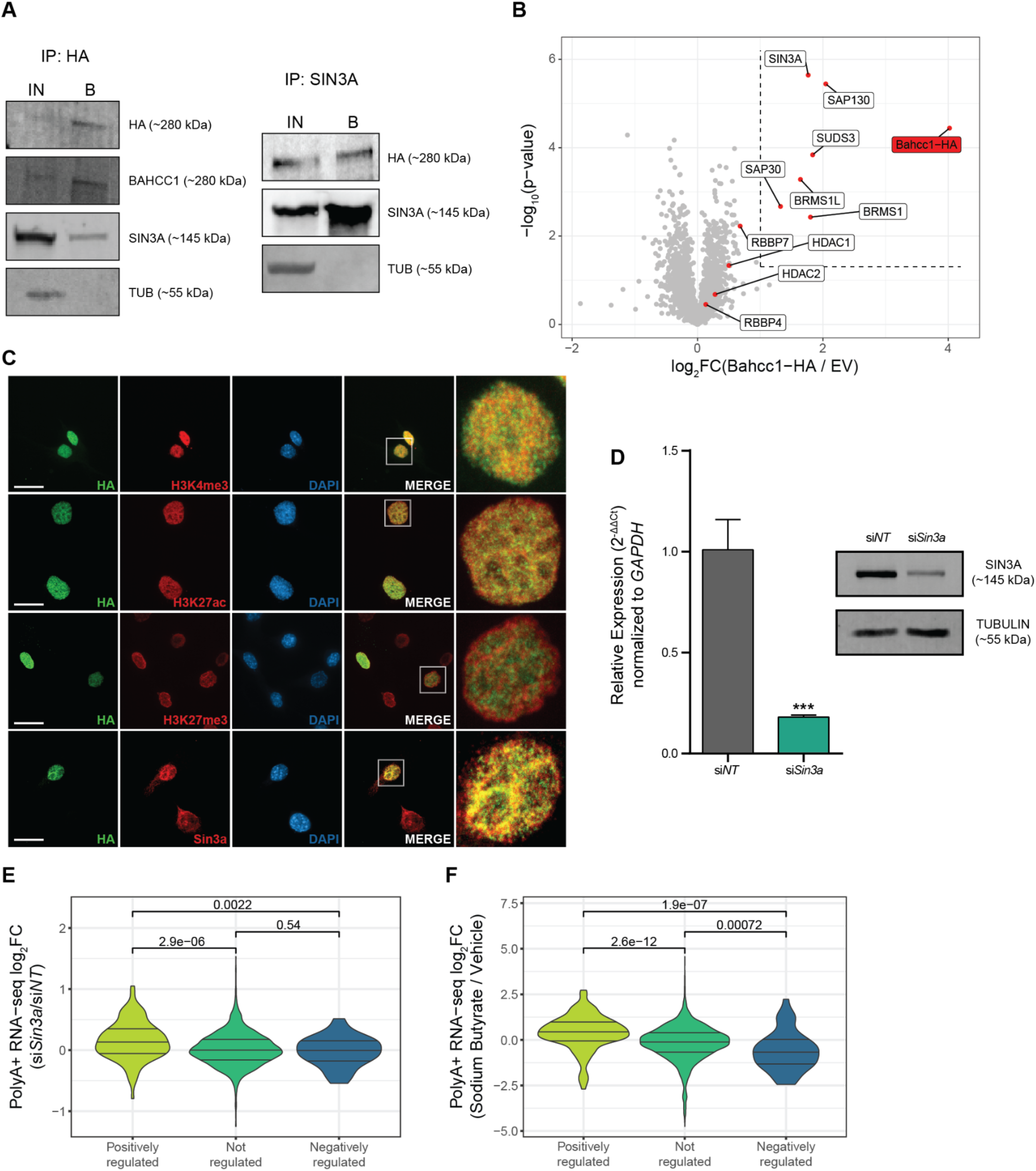
BAHCC1 associates with SIN3A and counteracts SIN3-HDAC activity. **(A)** Co-immunoprecipitation of BAHCC1 and SIN3A in HEK293T cells. **(B)** Volcano plot showing the magnitude and significance for the proteins identified by LC-MS after pulling down BAHCC1-HA. The dashed lines represent the p-value and log_2_FC thresholds, and all the core members of the SIN3A-HDAC complex have been highlighted. **(C)** Representative immunofluorescence images for BAHCC1 show overlap with H3K4me3, H3K27ac, and SIN3A, but not H3K27me3 (scale bar = 20 µm, 100X magnification). **(D)** RT-qPCR (left) and Western blot (right) upon *Sin3a* depletion with siRNAs to assess KD efficiency. **(E)** Changes in gene expression upon *Sin3a* KD for the positively and negatively regulated genes by BAHCC1. **(F)** Changes in gene expression after treating cells with sodium butyrate (pan-HDAC inhibitor) for the positively and negatively regulated genes by BAHCC1. All experiments were performed in n = 3 biological replicates, with the error bars in the barplots representing the standard deviation, p-value > .05 = ns; < 0.05 = *, < .01 = **; < .001 = *** (two-tailed Student’s t-test). In all violin plots, the three lines represent the median, first and third quartiles, and significance of the different comparisons was calculated by using a Mann-Whitney test.

We next tested if SIN3A also regulates the genes regulated by BAHCC1 by using siRNAs to deplete *Sin3a* in N2a cells (**Fig. 4D**). *Sin3a* KD promoted the expression of genes positively regulated by BAHCC1 (**Fig. 4E**). We then proceeded to KD other members of the SIN3-HDAC complex, specifically *Sin3b*, *Hdac1* and *Fam60a* (**Fig. S5A-C**). Interestingly, *Sin3b* KD had the opposite effect compared to *Sin3a*, with genes positively and negatively regulated by BAHCC1 being significantly reduced and induced by si*Sin3b*, respectively (**Fig. S5D**), which resonates with potentially opposing functions of the two proteins in other systems (see Discussion). *Hdac1* and *Fam60a* KD had no significant effect on the positively regulated genes but did affect the negatively regulated ones (**Fig. S5E-F**). To more efficiently inhibit HDAC activity, we treated N2a cells with sodium butyrate, a pan-HDAC inhibitor, resulting in large-scale gene expression changes (**Fig. S5G-H**). When considering genes regulated by BAHCC1, HDAC inhibition upregulated positively regulated genes and downregulated the negatively regulated ones (**Fig. 4F**), supporting the notion that these genes are regulated by BAHCC1 at least in part via histone acetylation.

These results are consistent with a model in which BAHCC1 is generally associated with H3K27ac-decorated regions, where its binding partners within the SIN3A complex are also recruited. At a subset of regions with stronger BAHCC1 association, BAHCC1 is positively regulating H3K27ac levels, possibly by inhibiting the HDAC activity of the SIN3A complex, which preferentially occupies promoter regions, but is also found at the BAHCC1-bound distal sites.

### BAHCC1 regulates histone acetylation and gene expression in primary cortical neurons

To test the relevance of the N2a cell line data in primary cells, we used the *Bahcc1*-targeting GapmeR to knock down *Bahcc1* in cultured primary mouse cortical neurons (**Fig. 5A**). Gapmer #1 was more effective in this system and thus was used in further experiments (**Fig. 5A**). We profiled BAHCC1, H3K27ac, H3K27me3, and H3K4me3 chromatin occupancy, changes in gene expression by RNA-seq, and changes in H3K27ac following GapmeRs free uptake. Consistently with the data in N2a cells, BAHCC1 binding was associated with active promoters (**Fig. 5B**), most correlated with H3K27ac histone mark (**Fig. 5C**), and knockdown by *Bahcc1* led to a decrease in H3K27ac levels in the peaks most strongly occupied by BAHCC1 (**Fig. 5D**). When considering changes in gene expression, *Bahcc1* KD led to significant changes in gene expression (**Fig. 5E**) and the up-regulated genes were enriched in terms related to synaptic and neurotransmitter signaling (**Fig. 5F**), whereas the down-regulated ones were enriched with genes acting in cell motility and neuronal development (**Fig. 5G**), consistent with the trends in the N2a data. Indeed, there was a weak yet significant correlation between changes in gene expression upon *Bahcc1* KD in N2a and in the primary cortical neurons (R=0.12, P<10^−15^). Among the 26 genes found to be significantly downregulated in both N2a and cortical neurons, some were previously linked to different functions in the nervous system, such as *Tnr* (36), *Gfra1* (GDNF receptor) (37, 38) and *Dkk3* (39). Of the 13 commonly upregulated genes, *Vgf* has been extensively linked to Alzheimer’s Disease (AD) and cognitive functions (40–43), *Atf5* to abnormal cortical development (44), *Emc1* to neurodevelopmental delay and cerebellar degeneration (45–48), and *Asns* to microcephaly and neuronal differentiation (49, 50). We analyzed different neuronal morphological features in *Bahcc1* KD primary cortical neurons (**Fig. 5H**) and found that cells with reduced *Bahcc1* expression exhibit reduced total neurite outgrowth (**Fig. 5I**, top) and number of processes (**Fig 5I**, top-center), which were shorter (**Fig. 5I**, center-bottom) and with fewer branches (**Fig. 5I**, bottom), suggesting that BAHCC1 activity is important for the development and branching of neurites.

**Figure 5.**
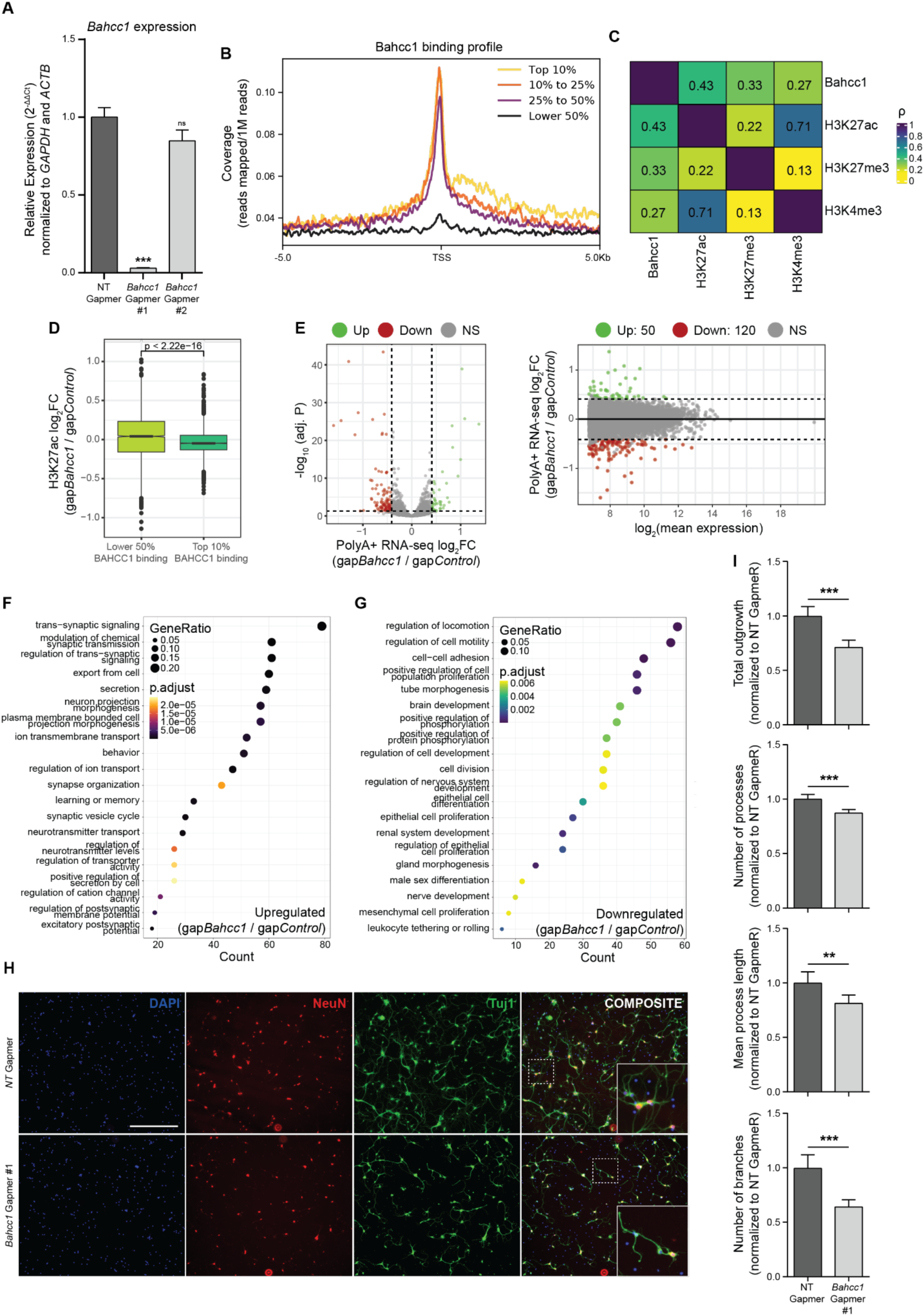
BAHCC1 affects gene expression and neurite outgrowth in primary cortical neurons. **(A)** RT-qPCR after 10 days of gymnotic transfer of *Bahcc1*-targeting GapmeRs in primary cortical neurons, to assess KD efficiency. **(B)** Metagene profile around TSS of the normalized BAHCC1 CUT&RUN signal in primary cortical neurons, stratified by gene expression levels. **(C)** Heatmap for the correlations of the CUT&RUN coverage between the examined proteins and histone modifications. Colors and numbers represent the Spearman’s correlation coefficients. **(D)** Changes in H3K27ac after *Bahcc1* KD at peaks with the lower 50% and top 10% average coverage of BAHCC1 CUT&RUN reads. **(E)** Vulcano (left) and MA (right) plots showing the gene expression changes in primary cortical neurons cultured with *Bahcc1*-targeting GapmeRs. **(F)** GO enrichment analysis of the upregulated genes following *Bahcc1* KD with GapmeRs. **(G)** Same as (F), but for the downregulated genes. **(H)** Representative images of primary cortical neurons 10 days after introduction of *Bahcc1*-targeting or control GapmeRs, stained with anti-Tuj1 and anti-NeuN antibodies. Scale bar = 200 µm, 10X magnification. **(I)** Normalized neurite outgrowth (top), number of processes (top-middle), average process length (middle-bottom) and number of branches (bottom) in primary cortical neurons cultivated in the presence of either *Bahcc1*-targeting or control GapmeRs. All experiments were performed in n = 3 biological replicates, except for neuronal morphological analyses which were done with n = 6, with the error bars in the barplots representing the standard deviation. P > .05 = ns; < 0.05 = *, < .01 = **; < .001 = *** (two-tailed Student’s t-test). For the boxplot in (D), the thick line, edges of the box, and whiskers represent the median, first and third quartiles, and the upper and lower 1.5 interquartile ranges (IQRs), respectively. Outliers (observations outside the 1.5 IQRs) are drawn as single points, and the significance was calculated by a two-sided Wilcoxon rank-sum test. Genes were considered to be differentially expressed when adj. P < 0.05 and |log_2_FC| > .41 (corresponding to a change of 33%).

Taken together, our data highlight a role for BAHCC1 in primary cortical neurons, where it affects the expression of several genes important for neuronal biology. These changes are eventually translated into an impaired growth of neurites, both in terms of length and number of branches, suggesting that BAHCC1 plays an important role in neuronal function.

## Discussion

The Sin3 complex, first discovered in yeast in 1987 (51, 52) is centered around the SIN3A/SIN3B scaffold proteins in vertebrates.It is generally thought to be repressive, as it contains histone deacetylases, traditionally associated with gene silencing, a function well established in yeast (53). However, genome-wide occupancy studies in mammalian cells have found the Sin3 complex to be associated with many active promoters and enhancers (13–15). The diversity of complex isoforms and substoichiometric components has been suggested to underpin this apparent discrepancy (14). BAHCC1 is a strong candidate for a possible modulator of SIN3A activity (**Fig. 6A** and **6B**). BAHCC1 levels are apparently substoichiometric to SIN3A. In HEK293T cells there are an estimated 1.6×10^3^ copies of BAHCC1 compared to 1.9×10^5^ estimated copies of SIN3A and at least 3×10^4^ copies of the other SIN3A core components HDAC1, SAP30, FAM60A, RBBP4, RBBP7, and ING2 (https://opencell.czbiohub.org/) (**Fig. 6A**). TNRC18, a paralog of BAHCC1 is also typically more abundant than BAHCC1. Similarly, while *Bahcc1* mRNA is expressed in all mouse tissues and developmental stages, its levels are generally modest, with a slight enrichment in the developing brain (21). Also, the phenotype of loss of BAHCC1 is less dramatic than that of Sin3. *Sin3a*^−/−^ mouse embryos die before day E6.5 (54), *Rbbp4*^−/−^ die before E7.5 (55) whereas *Bahcc1*^−/−^ die shortly after birth (27). All these suggest that although BAHCC1 is consistently recovered in pulldowns of Sin3 complex components in various systems (**Fig. 4A**, 4B and (13, 16, 23–26)), it is not a core component of this complex but rather plays an auxiliary, and likely regulatory role.

**Figure 6.**
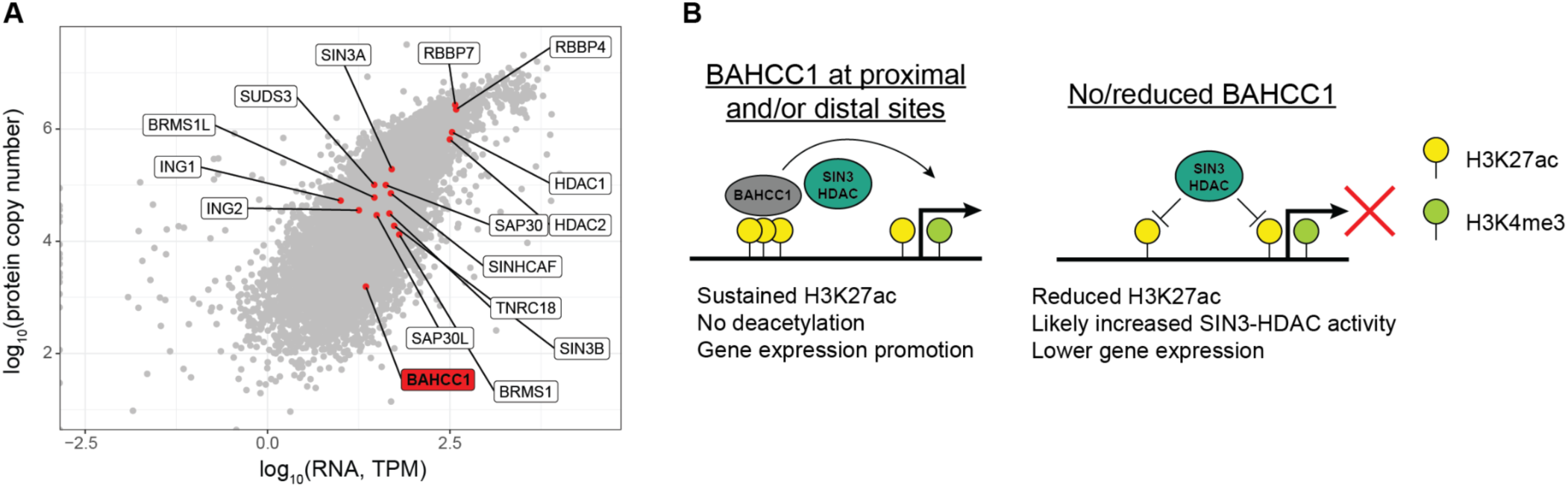
BAHCC1 acts substoichiometrically to modulate SIN3-HDAC. **(A)** RNA and corresponding protein levels for Sin3 components and BAHCC1 in HEK293T cells (data from https://opencell.czbiohub.org/). **(B)** Proposed model for BAHCC1 modulatory activity on SIN3-HDAC.

Interestingly, we find that genes positively regulated by BAHCC1 are up-regulated by KD of *Sin3a*, but also significantly down-regulated by loss of *Sin3b* (**Fig. 4D** and **S5D**). Consistently, perturbations of SIN3A and SIN3B described in the literature result in different phenotypes (56, 57) and even opposing effects on breast cancer metastasis (58), in line with the differences between their patterns of protein-protein interactions (59). Still, the relationship between the two proteins and the complexes they scaffold is largely unknown. Intriguingly, we did not find any correlation between the effects of *Sin3a* and *Sin3b* knockdowns (R=0.02; P=0.09), so we suggest that BAHCC1 target sites may define regions with specific opposing effects of the two proteins, that can be further elucidated in future studies. Moreover, BAHCC1 pulldown in HEK293T cells specifically retrieved SIN3A but not SIN3B (**Fig. 4B**), further suggesting that BAHCC1 activity is primarily linked to the former and mostly affects the activity of the SIN3A-containing HDAC complexes.

A previous study has linked BAHCC1 primarily to H3K27me3 chromatin marks(16), whereas we do not see any evidence that BAHCC1 co-localizes with H3K27me3 in our CUT&RUN or immunofluorescence data. A possible explanation is that we’re generally examining the full-length BAHCC1 protein, whereas (16) mostly studied just its BAH domain that corresponds to <5% of the full-length protein. Notably, some of our results are consistent with those of (16). In both studies, KD of HDAC1 and SIN3A components lead to the de-repression of BAHCC1-associated genes. In leukemic cells, the model proposed by Fan et al. was that BAHCC1 recruits SIN3A to regions demarcated by H3K27me3. In our model, BAHCC1-bound regions respond to HDAC inhibition because they are also SIN3A targets, with BAHCC1 likely moderating the efficacy of their repression.

Functionally, we find that loss of BAHCC1 leads to decreased expression of genes enabling cellular migration, and a reduction in N2a motility in transwell migration and wound healing assays. The ability of neuronal progenitors to migrate is a key component of their biology (60), and a role for BAHCC1 in enabling this migration may explain its preferential expression in the developing over the adult brain (21). Consistently with this model, perturbations of BAHCC1 reduced cellular motility and invasiveness in melanoma cells as well (17), and conversely, perturbation of the *SIN3A, FAM60A* and *SUDS3* Sin3 components led to increasing invasiveness and enhanced cell migration (25, 58, 61, 62). In primary mouse cortical neurons harvested from the developing (E16.5/E17) brain, *Bahcc1* depletion caused decreased total neurite outgrowth, with neurons exhibiting fewer and shorter processes, as well as reduced number of branches (**Fig. 5I**). This is consistent with the observed transcriptomic changes that mostly encompass genes involved in cell motility, neuronal morphogenesis, synapse development and neurotransmitter signaling. Notably, loss-of-function of some of the BAHCC1-regulated genes are associated with relevant phenotypes, such as GFRα1 (38), Dkk3 (39), Vgf (63), and Atf5 (44).

We previously studied *Bahcc1* in the context of neuronal differentiation of mESCs (21). In that system genetic manipulation of *Bahcc1* led to massive cell death upon neuronal differentiation, a more drastic phenotype than the one we observe here in more mature cells, possibly because of the only partial knockdown we can achieve with GapmeRs. In our previous study we found that *Bahcc1* perturbation, similarly to the perturbation of the *Reno1* long noncoding RNA, that is transcribed from a region ∼45 Kb upstream of *Bahcc1*, led to a strong differentiation defect, and loss of DNA accessibility at regions marked with H3K4me3. These results are consistent with the model where BAHCC1 is affecting regions that positively regulate gene expression. Early neuronal differentiation entails a very complex series of chromatin remodeling, likely much more than N2a cells before or after RA-directed differentiation. N2a cells express *Reno1*, but only at very low levels of ∼1 FPKM compared to 3–4 FPKM in the early neurons (21), so our attempts to perturb *Reno1* in this system did not lead to consistent results. Interestingly, over-expression of *Reno1* in this system, which leads to expression of *Reno1* far beyond its physiological levels (**Fig. S6A**), down-regulates the genes positively regulated by Bahcc1 (**Fig. S6B**) and is significantly anti-correlated with BAHCC1 OE in N2a cells (R=–0.42 P<10^−15^, **Fig. S6C**), in an apparent contrast to the co-expression and similar functions of the two genes in mESCs. Future studies of *Reno1* in a system with higher levels, such as the developing brain or early NPCs, may shed further light on its roles.

Transcription is regulated by a complex interplay between a large number of different players that act in part by modifying the chromatin and in part by recruiting or repelling different components of the transcription initiation or elongation/pause-release machinery. This process is likely most complex in the lineages where cell division is limited, and where there is a large diversity of cell types, such as the developing and the mature nervous system. The preferential expression of *Bahcc1* in the brain, and in particular in the developing brain (21), and the neuronal phenotypes in the *Bahcc1*^−/−^ mice suggest that it plays a prominent role in sculpting gene expression in the developing and potentially the mature brain.

## Materials and Methods

### Cell culture and transfection

Neuro2a (N2a) cells were cultured in DMEM supplemented with 4.5 g/L of glucose, 4 mM L-Glutamine, 10% FBS, 1% PenStrep solution, grown in incubators maintained at 37°C with 5% CO_2_, and were routinely passaged by gentle dissociation using a Trypsin (0.05%) – EDTA (0.02%) solution. DNA plasmid transfections were performed by using Lipofectamine 3000 (ThermoFisher Scientific), according to the respective protocols. For each transfection, the optimal DNA concentration and amount of transfection reagents were empirically determined to minimize cellular stress and maximize efficiency, to eventually ensure clean readouts. To induce the expression of doxycycline-inducible plasmids, cells were cultured in the presence of doxycycline at a final concentration of 2 µg/µL. SiRNAs were transfected using DharmaFECT reagents (DharmaFECT 2), at a final concentration of 25 µM concordantly to the manufacturer’s protocol. GapmeRs were transfected at the final concentration of 50 µM with Lipofectamine 3000, following the manufacturer’s guidelines. In all cases, cells were plated the day before the transfection in an antibiotic-free medium which was subsequently changed prior to adding the transfection mix, and again after 24h to minimize cell death. Finally, transfections were carried out for 72h, unless stated otherwise. The complete list with the catalog numbers and sequences of both siRNAs and LNA/GapmeRs is reported in **Table 1**.

### Plasmid construction

The sequence corresponding to the cDNA of mouse *Bahcc1* (ENSMUST00000118987) minus the STOP codon was synthetized by Hylabs and cloned inside the pLIX_402 vector (gift from David Root, Addgene #41384) by PCR cloning via 5’ NheI and 3’ AgeI sites by recombination. To generate a plasmid encoding a full-length *Bahcc1* without the BAH domain (Bahcc1^ΔBAH^-HA), we digested the plasmid with AgeI to remove the last 918bp of the *Bahcc1* CDS. All plasmids were sent for Sanger and whole plasmid (Plasmidsaurus) sequencing prior to use.

### Primary neuronal culture and morphological analysis

Embryonic cortices (embryonic day 16.5–17) were dissected from 5–6 embryos of mixed sex and pooled for a single replicate culture. Trypsin (0.25%) was used to dissociate tissue through a 10 minutes incubation at 37°C. Digestion was terminated with the addition of ovomucoid (trypsin inhibitor, Worthington). Digested tissue was gently triturated through a P1000 pipette to release cells and passed through a 40 µm filter. After trituration, the cell suspension was transferred to a 15 mL Falcon tube and centrifuged for 5 minutes at RT / 154×g. After centrifugation, the cloudy supernatant was carefully removed with a pipette and discarded. The pellet was resuspended in 10 mL DMEM medium and added to uncoated 90 mm Petri dishes. The plates were incubated at 37°C, 5% CO2 humidified incubator for 30 minutes. During this incubation, glial cells and other unwanted debris adhere to the bottom of the plate while cortical cells are retained in the media and recover from the trituration. The supernatant was carefully removed, without disturbing attached glial cells and unwanted cell debris at the bottom of the plate, and moved into a new 15 mL Falcon tube using a pipette. The collected supernatant was then centrifuged for 5 minutes at RT / 154×g. Neurons were plated onto cell culture dishes pre-coated overnight with poly-d-lysine (20 µg/mL) and laminin (4 µg/mL) and were grown in neurobasal medium (Gibco) containing B27 supplement (2%), 1% PenStrep solution and glutaMAX (1 mM). Neurons were grown in incubators maintained at 37°C with a CO2 concentration of 5%. GapmeRs for unassisted uptake were added directly to the culture medium at a final concentration of 500 nM, and the neurons were cultured for a total of 10 days. Neuronal images were acquired at X10 magnification on an ImageXpress Micro (Molecular Devices) automated microscopy system and quantified using WIS-Neuromath (64). The parameters reported include total outgrowth, defined as the sum of lengths of all processes and branches per cell,, the number of processes, the average process length and the number of branching points.

### Sodium Butyrate and Retinoic Acid treatments

Sodium Butyrate (Sigma) was diluted in sterile H_2_O at a stock concentration of 1M and added to the cell culture media at a final concentration of 5µM. As a control, cells were supplemented with an equal volume of H_2_O. All treatments were performed for 24h in order to minimize indirect effects due to the extensive inhibition of deacetylation. Retinoic Acid (Sigma) was diluted in DMSO at a stock concentration of 0.01M, and added to a complete media with reduced (1%) serum concentration at a final concentration of 10µM, with equal volumes of DMSO serving as vehicle controls. After three days of treatment, cells were transfected with GapmeRs as previously described, and incubated for three additional days in the retinoic acid medium.

### Gene expression analysis

RNA was extracted by using TRI-Reagent, according to the manufacturer’s instructions. Briefly, cells were washed once with 1X PBS and the proper amount of TRI was added directly to the wells. After complete lysis, 200 µL of chloroform for each mL of TRI was added and the samples were centrifuged at 4°C / 15000 RPM for 15 minutes, and the resulting upper aqueous phase was isolated. RNA was then precipitated by adding 500 µL of 2-propanol for each mL of TRI, incubated at room temperature for 15 minutes while rotating and centrifuged for 15 minutes at 4°C / 15000 RPM. The RNA pellet was then washed once with 1 mL of 75% ethanol and dried. RNA was resuspended in DNase/RNase-free H_2_O. Baseline-ZERO™ DNase (Biosearch Technologies) was applied to 1-2 µg of total RNA, following the manufacturer’s recommendations and incubating for 1 hour at 37°C. DNase-treated or untreated RNA was retrotranscribed using the qScript™ Flex cDNA Synthesis Kit (Quanta Bio), with either oligo-dT or a 1:1 combination of oligo-dT and random hexamers, following standard conditions as suggested by the manufacturer. The resulting cDNA was then diluted 1:5-1:10 with DNase/RNase-free H_2_O and used as input for Quantitative Real-Time PCR using the Fast SYBR™ Green Master Mix (Applied Biosystems). Each reaction (5 µL Fast SYBR™ Green Master Mix, 1 µL diluted cDNA, 3.75 µL DNase/RNase-free H_2_O and 0.25 µL of 10 µM Forward + Reverse primers) was carried out in triplicate and amplified using the following PCR program: 20’’ at 95°C, 40 cycles by denaturing for 1” at 95°C followed by annealing/extension for 20’’ at 60°C. Results were analyzed using the ΔΔCt method. The complete list of primers and their sequences is available in **Table 2**.

### Protein extraction and Western Blot

Proteins were extracted using RIPA Buffer (150mM NaCl, 1% NP-40, 0.5% Deoxycholate, 0.1% SDS, 50mM TRIS pH 8, 1mM DTT), freshly supplemented with EDTA-free Protease Inhibitor Cocktail (APExBIO)) and 1mM DTT. Briefly, cells were detached with a Trypsin (0.05%) – EDTA (0.02%) solution, pelleted by centrifugation, resuspended in ice-cold RIPA buffer, and incubated on ice for 20 minutes with occasional vortexing. The lysates were then centrifuged at 15,000 rpm for 15 minutes at 4°C, and the supernatants were collected, quantified by Bradford Assay, and stored at -80°C. For Western Blots, an equal amount of proteins was loaded on a polyacrylamide gel and run in SDS Running Buffer. Proteins were then transferred to a pre-activated 0.2 or 0.45µm PVDF membrane in a cold Tris-Glycine Buffer supplemented with 20% methanol inside ice, with a constant current of 0.30A for 2 hours. Blocking was done in 5% milk in 1X PBS-0.1% Tween 20 for at least 1h, after which the membranes were incubated O/N at 4°C with the primary antibodies diluted in 5% milk in 1X PBS-0.1% Tween 20. The following day, the membranes were washed three times with 1X PBS-0.1% Tween 20, incubated for 2h at RT with the appropriate secondary antibodies in 5% milk in 1X PBS-0.1% Tween 20, and washed again three times prior to image acquisition. The complete list of primary and secondary antibodies and relative dilutions is available in **Table 3**.

### Co-immunoprecipitation

A total of 5-10×10^6^ cells were detached with a Trypsin (0.05%) – EDTA (0.02%) solution, pelleted by centrifugation and frozen at -80°C for at least 1 hour. The pellets were then thawed, resuspended in 450 µL of ice-cold lysis buffer (1% Triton X-100, 50 mM Tris-HCl pH 7.4, 150 mM NaCl) freshly supplemented with EDTA-free Protease Inhibitor Cocktail (APExBIO)) and incubated on ice for 20 minutes with occasional vortexing. For each sample, 50 µL of Protein A/G paramagnetic beads were washed three times with ice-cold lysis buffer, and resuspended in 250 µL of ice-cold lysis buffer. The lysates were then centrifuged at 15,000 rpm for 10 minutes at 4°C, the supernatants were collected into a new tube and pre-cleared with 50 µL of unconjugated beads for ∼30 minutes at RT, while gently rotating. The rest of the beads were incubated with 5 µg of antibody for ∼1h at RT with gentle rotation, after which they were washed again three times with ice-cold lysis buffer to remove any excess antibody, and finally incubated with the pre-cleared protein lysates overnight at 4°C, while gently rotating. The following day, the beads were washed three times with ice-cold lysis buffer to remove unbound proteins. For Western blot analysis, the beads were eluted directly in a loading buffer and incubated for 5 minutes at 95°C, and the eluate was either stored at –80°C or directly loaded on a gel. The complete list of antibodies and relative dilutions is available in **Table 3**.

### Sample preparation, liquid chromatography mass spectrometry and data processing

The samples were subjected to tryptic digestion using an S-trap(65). The resulting peptides were analyzed using nanoflow liquid chromatography (Acquity M-Class) coupled to high resolution, high mass accuracy mass spectrometry (Q Exactive HFX). Each sample was analyzed on the instrument separately in a random order in discovery mode. Raw data was processed with MetaMorpheus v1.0.2. The data was searched against the human Uniprot proteome database (xml version including known PTMs) appended with the BAHCC1-HA sequence, common lab protein contaminants and the following modifications: Carbamidomethylation of C as a fixed modification and oxidation of M as a variable one. Quantification was performed using the embedded FlashLFQ(66) and protein inference(67) algorithms. The LFQ (Label-Free Quantification) intensities were calculated and used for further calculations using Perseus v1.6.2.3. Decoy hits were filtered out. The LFQ intensities were log transformed and only proteins that had at least 2 valid values in at least one experimental group were kept. The remaining missing values were imputed. A student’s t-test was performed to identify the proteins that are differentially expressed.

### Cleavage Under Targets and Release Using Nuclease (CUT&RUN)

CUT&RUN reactions were performed following the V3 of the protocol (28), with minor modifications. An equal number of cells (up to 500,000 for each sample/replicate) was harvested using a Trypsin (0.05%) – EDTA (0.02%) solution and washed once with 1X PBS at room temperature. After three washes with Wash Buffer (20mM HEPES-NaOH pH 7.5, 150mM NaCl, 0.5mM Spermidine supplemented with EDTA-free Protease Inhibitor Cocktail (APExBIO)) at RT, cells were resuspended in 1 mL Wash Buffer and incubated while gently rotating with 20 µL of Concanavalin A-coated magnetic beads (EpiCypher), which have been previously washed, activated and resuspended with Binding Buffer (20mM HEPES-NaOH pH 7.5, 10mM KCl, 1mM CaCl_2_, 1mM MnCl_2_). After 10’, the buffer was removed, and the beads were resuspended in 150 µL of Antibody Buffer (Wash Buffer with 0.1% Digitonin (Sigma) and 2mM EDTA pH 8) containing the proper dilution of the antibody of interest and left gently rotating overnight with a 180° angle at 4°C. On the next day, cells were washed two times with ice-cold Dig-Wash Buffer (Wash Buffer with 0.1% Digitonin), resuspended in 150 µL of Dig-Wash Buffer supplemented with 1 µL of custom-made pA/G-MNase every mL and left rotating for 1h at 4°C. After that, cells were washed again two times with Dig-Wash Buffer, resuspended in 100 µL of Dig-Wash Buffer, and placed into a thermoblock sitting on ice. To initiate the cleavage reaction, 2 µL of a freshly diluted (from a 1M stock) solution of 100mM CaCl_2_ was added to each tube, and the tubes were left at 0°C for 30 minutes. To halt the reaction, 100 µL of 2X STOP Buffer (340mM NaCl, 20mM EDTA pH 8, 4mM EGTA pH 8, 0.05% Digitonin, 100 µg/mL RNAse A, 50 µg/mL Glycogen) were added to each tube, and cleaved DNA fragments were released by incubating the samples for 30’ at 37°C. The tubes were then centrifuged for 5 minutes at 4°C / 16,000g and the supernatant was collected. Finally, the DNA was purified by standard Phenol/Chloroform extraction using the 5PRIME Phase Lock Gel Heavy tubes (QuantaBio), and the success of the CUT&RUN reaction was assessed by running the positive control (H3K27me3 or H3K4me3) on a Tapestation (Agilent Technologies) using a High Sensitivity D1000 ScreenTape (Agilent Technologies). When using the H3K27ac, SIN3A, and BAHCC1 antibodies, all buffers were supplemented with 20mM of sodium butyrate (Sigma) to prevent the loss of acetylated residues. The complete list of primary and secondary antibodies and relative dilutions is available in **Table 3**.

### RNA-seq and CUT&RUN Library Preparation

RNA-seq libraries were prepared using the CORALL mRNA-Seq Kit V2 (Lexogen), following the manufacturer’s instructions. Prior to library generation, 1 µg of total RNA was PolyA-enriched using the Poly(A) RNA Selection Kit V1.5 (Lexogen). To prepare CUT&RUN libraries, we followed the original protocol (28), with slight modifications. Briefly, to 30 µL of purified CUT&RUN DNA we added 10 µL of a 4X End Repair and A-Tailing (ERA) Buffer, consisting of 4 µL of 10X T4 DNA Ligase Buffer (NEB), 2 µL 10mM dNTPs mix, 1 µL 10mM ATP, 2 µL of 50% PEG 4000 (ThermoFisher Scientific), 0.6 µL PNK (NEB), 0.1 µL T4 DNA Polymerase (NEB) and 0.3 µL of Taq Polymerase (homemade). Samples were incubated in a thermocycler pre-heated at 12°C with the following program: 15 minutes at 12 °C, 15 minutes at 37°C, 45 minutes at 58°C, hold at 8°C. Barcoding was performed by adding to each sample 54 µL of 2X DNA Quick Ligase Buffer (NEB), 4 µL of DNA Quick Ligase (NEB) and 10 µL of unique Y-shaped adaptors (final concentration of 0.075 µM). The samples were incubated for 20’ in a pre-heated thermocycler at 20°C, after which 1 µL of 20% SDS and 2 µL of 20 mg/mL Proteinase K were added and the temperature was raised to 37°C for 1h. Following purification with 1X SPRI beads (GE Healthcare) and a second round with 1.2X HXP Buffer (20% PEG 8000, 2.5M NaCl), libraries were amplified by a PCR reaction composed of 25 µL 2X KAPA HiFi HotStart ReadyMix (Roche), 2 µL of Primer Mix (Illumina TruSeq Universal and Illumina P7 standard primers, at 50 µM each) and 23 µL of purified DNA from the previous step, with a PCR program consisting of 45’’ at 98°C, 14 cycles of 15’’ at 98°C and 10’’ at 60°C, with a final extension of 1 minute at 72°C. Amplified libraries were subsequently purified with 1.1X SPRI beads and then again with 1.2X HXP Buffer. The libraries were then quality-checked by both dsDNA Qubit (Thermo Fisher Scientific) and Tapestation (Agilent Technologies). All libraries were sequenced on an Illumina NovaSeq 6000 or Novaseq X instrument, aiming for either 10M (CUT&RUN) or 15M (polyA+ RNA-seq) reads per sample.

### RNA-seq data analysis

Raw FASTQ files were processed by an adapted Lexogen CORALL analysis pipeline script (https://github.com/Lexogen-Tools/corall_analysis). Briefly, adaptors were trimmed by using Cutadapt(68), and the resulting trimmed FASTQ files were used as input for mapping to the mm10 mouse genome with STAR aligner(69), using the ‘--quantMode GeneCounts’ option in order to count the number of reads mapping to each feature provided by an appropriate GTF file. To call for differential expression, we used DESeq2 (31), considering a gene to be differentially expressed when adjusted P < 0.05 and absolute log_2_FC > 0.41 (∼33% increase or decrease). Prior to any subsequent analysis, we filtered out poorly expressed genes, pseudogenes, and genes with an exonic length < 200nt. To perform GO enrichment analysis and GSEA, we used the Clusterprofiler R package (70) by considering a term to be enriched when the adj. P < 0.05. Whenever several redundant terms were found, we employed the *simplify* function with a cutoff=0.7 to reduce the redundancy.

### CUT&RUN data analysis

Raw FASTQ files were aligned to the mm10 reference genome with Bowtie2(71), using the -p 8, -X 2000 and --no-unal options. BAHCC1 and SIN3A peaks were called using the MACS2 callpeak function(72), whereas for H3K27ac, H3K4me3 and H3K27ac we instead used epic2(73). Peak quantification and differential expression were performed using the HOMER tool suite (74), using the *annotatePeaks.pl* and *getDiffExpression.pl* functions, respectively. To compute the correlation between individual bigWig files and to plot the metagene profiles and coverage heatmaps, we employed *bigWigCorrelate* and deepTools(75), respectively.

### Immunostaining

Cells were plated and transfected on either sterile glass coverslips or 8-well chambers (Ibidi). Cells were first washed once with 1X PBS, and then fixed with cold 4% PFA in 1X PBS for 15 minutes at RT. After two washes with 1X PBS for 5 minutes each, cells were permeabilized by incubating them with 5% horse serum, 1 mg/mL BSA, 0.1% Triton-X100 in 1X PBS for 30 minutes at RT. Following three washes with 1X PBS at RT, cells were incubated O/N at 4°C with the primary antibodies in blocking solution (5% horse serum, 1 mg/mL BSA in 1X PBS). The day after, cells were washed three times with 1X PBS and incubated for 2h at RT with the secondary antibodies in a blocking solution, protected from light. Lastly, cells were washed three times with 1X PBS, stained with DAPI, and mounted with ProLong™ Glass Antifade Mountant (Invitrogen). Slides were left curing in the dark O/N at RT, and eventually imaged with a Zeiss Spinning Disk confocal microscope. The complete list of primary and secondary antibodies and relative dilutions is available in **Table 3**.

### Wound Healing Assay and Analysis

To perform the wound healing assay, we employed the µ-Plate 24 Well system (Ibidi) in order to increase consistency between scratches and standardize the experimental conditions. Moreover, to avoid the need to detach and plate again cells that already underwent transfection, we introduced the targeting and control LNAs via reverse transfection, employing the same conditions as in the regular settings and diluting the transfection mixes directly into the cell mixtures while seeding. A total of six replicates for each condition were seeded and transfected. After reaching confluence overnight, the culture inserts were gently removed with sterile forceps and the wells were washed with 1X PBS to remove any cell debris. Brightfield images were taken using an EVOS Cell Imaging System (Thermo Fisher Scientific) at different time points. Data analysis was then performed using the Wound_healing_size_tool (76) plugin for ImageJ, which calculates the total (µm^2^) and percentage (%) of the wounded area. From these data, we calculated the the percentage of wound closure as follows:

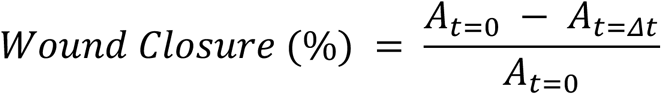

With A_t=0_ and A_t=Δt_ being the wounded area (in µm^2^) at the beginning and after time t, respectively. The raw and normalized (on time 0) area were also used as parameters, given the highly standardized experimental setting.

### Transwell assay

To measure cell migration, we performed a transwell invasion/migration assay using transwell inserts with 8.0 µm pores (Corning). Briefly, cells were plated and reverse transfected as described above in low serum (1%) media, on the upper part of each insert. In the bottom part, we added regular media (10% serum) in order to attract the cells. After 24 hours, non-migrated cells were scraped off using sterile cotton swabs, after which the wells were washed three times with 1X PBS. The migrated cells on the lower part were then fixed with ice-cold 4% PFA for 15 minutes at room temperature, washed twice with 1X PBS and stained with crystal violet for 10 minutes, after which they were washed two more times with 1X PBS. Crystal violet was then extracted by adding a 33% Acetic Acid solution for 10 minutes, while gently rocking the plate. Finally, optical density at 595 nm was measured using a Cytation 5 microplate reader (BioTek).

## Data availability

All the RNA-seq and CUT&RUN data are available at the GEO database accession GSE281267. The mass spectrometry data have been deposited in the PRIDE database under the accession PXD058811.

---

**Supplementary Figure 1.**
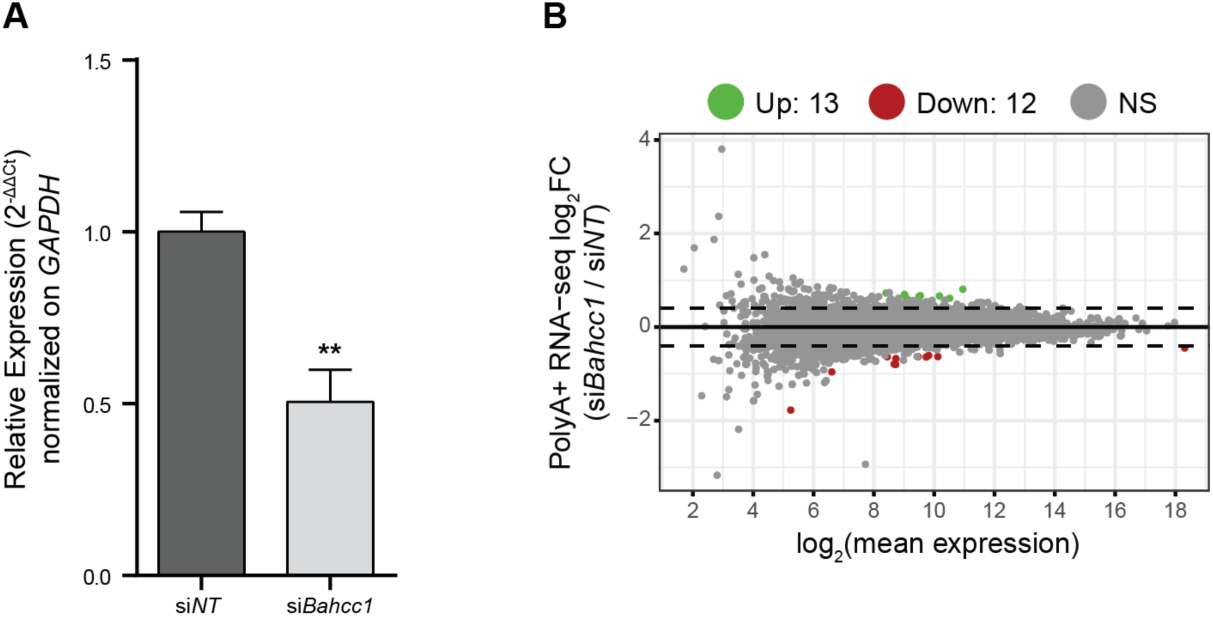
BAHCC1 KD with siRNAs in N2a is only modestly efficient. **(A)** RT-qPCR upon *Bahcc1* depletion with siRNAs to assess KD efficiency. **(B)** MA plot showing the genome-wide gene expression changes after *Bahcc1* KD with siRNAs. All experiments were performed in n = 3 biological replicates, with the error bars in the barplots representing the standard deviation, P > .05 = ns; < 0.05 = *, < .01 = **; < .001 = *** (two-tailed Student’s t-test). Genes were considered to be differentially expressed when adj. P-value < 0.05 and |log_2_FC| > 0.41 (corresponding to a change of 33%).

**Supplementary Figure 2.**
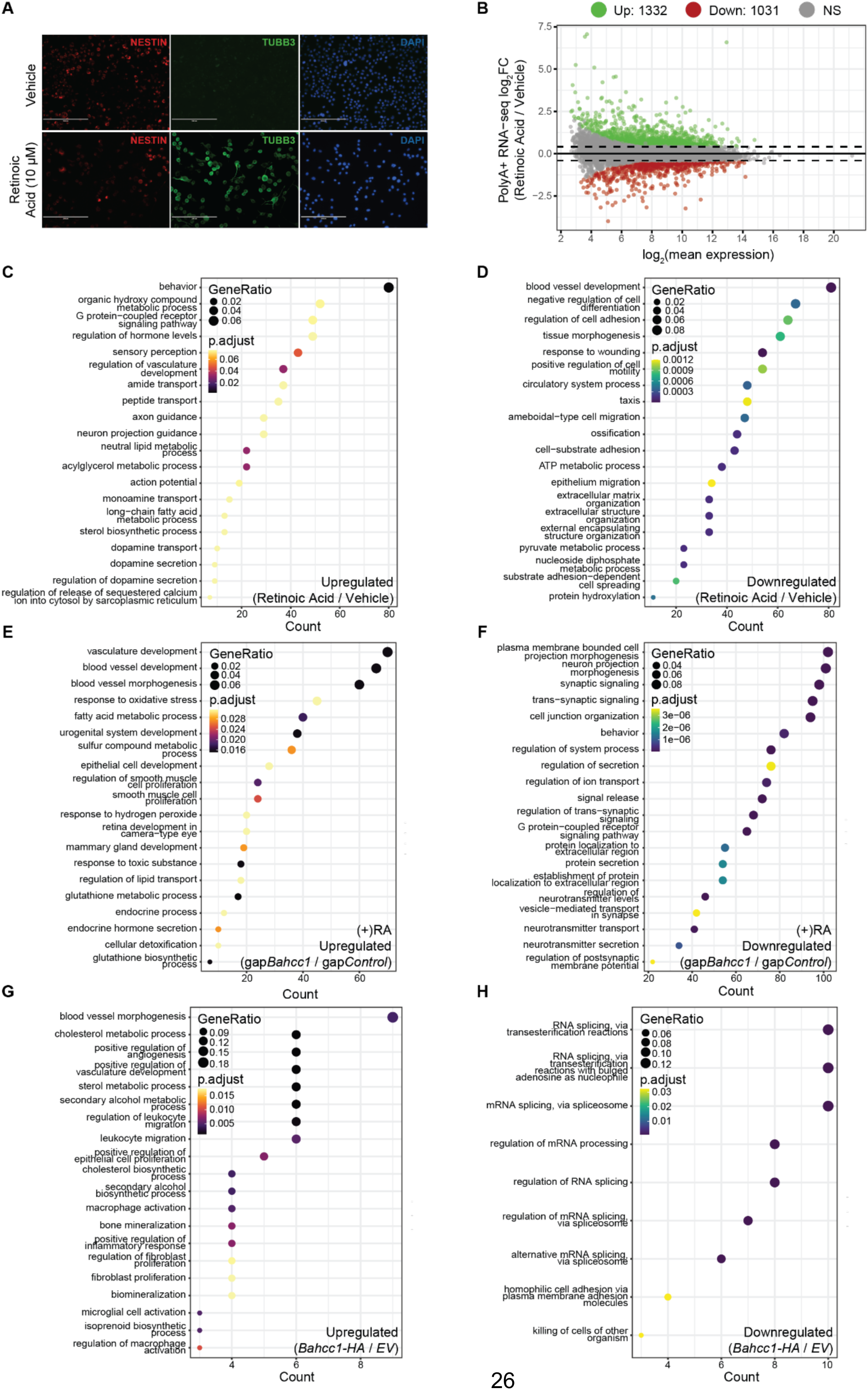
BAHCC1 affects gene expression in both undifferentiated and differentiated N2a cells. **(A)** Representative images of N2a cells after three days of RA treatment, staining for markers of neural progenitors (NESTIN) and mature neurons (TUBB3) by immunofluorescence (10X magnification). **(B)** MA plot showing the genome-wide gene expression changes after three days of differentiation with retinoic acid. **(C)** GO enrichment analysis of the genes upregulated by RA treatment. **(D)** Same as in (C), but for the downregulated genes. **(E)** GO enrichment analysis of the upregulated genes after *Bahcc1* KD with GapmeRs in RA-treated N2a cells. **(F)** Same as in (E), but for the downregulated genes. **(G)** GO enrichment analysis of the genes upregulated after *Bahcc1* OE. **(H)** Same as in (G), but for the downregulated genes. All experiments were performed in n = 3 biological replicates. Genes were considered to be differentially expressed when adj. P-value < 0.05 and |log_2_FC| > 0.41 (corresponding to a change of 33%).

**Supplementary Figure 3.**
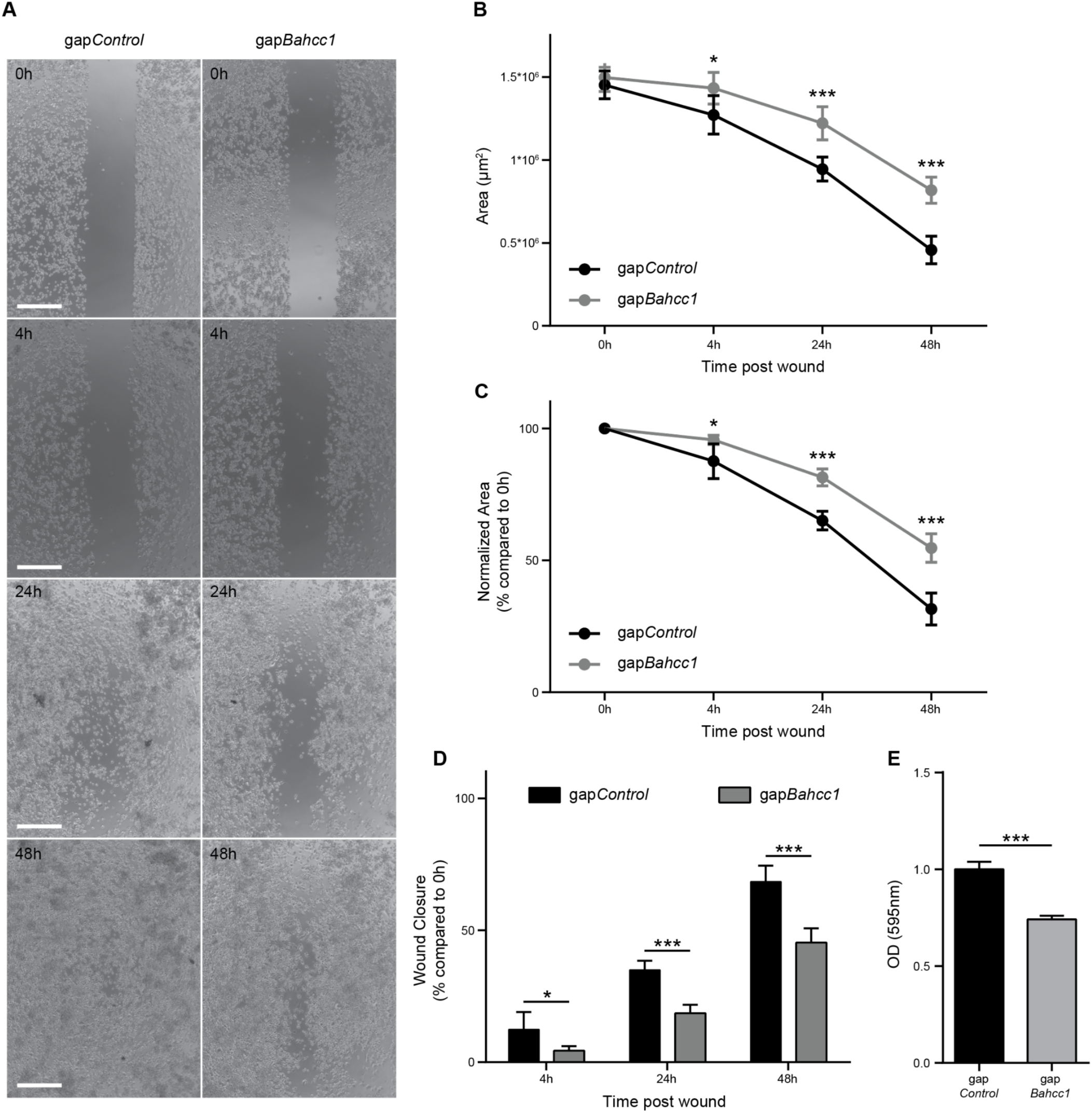
Depletion of BAHCC1 affects N2a cells invasion and migration. **(A)** Representative brightfield images during the time-course of the wound healing experiment at the reported time points (scale bars = 500 µm, 4X magnification). **(B)** Absolute wounded area at the reported time points. **(C)** Normalized wounded area (compared to 0h) at the reported time points. **(D)** Wound closure percentage (compared to 0h) at the reported time points. **(E)** Crystal Violet absorbance of migrated cells. All experiments were performed in n = 3 (transwell assay) or n = 6 (wound healing assay) biological replicates, with the error bars in the barplots representing the standard deviation, P > .05 = ns; < 0.05 = *, < .01 = **; < .001 = *** (two-tailed Student’s t-test).

**Supplementary Figure 4.**
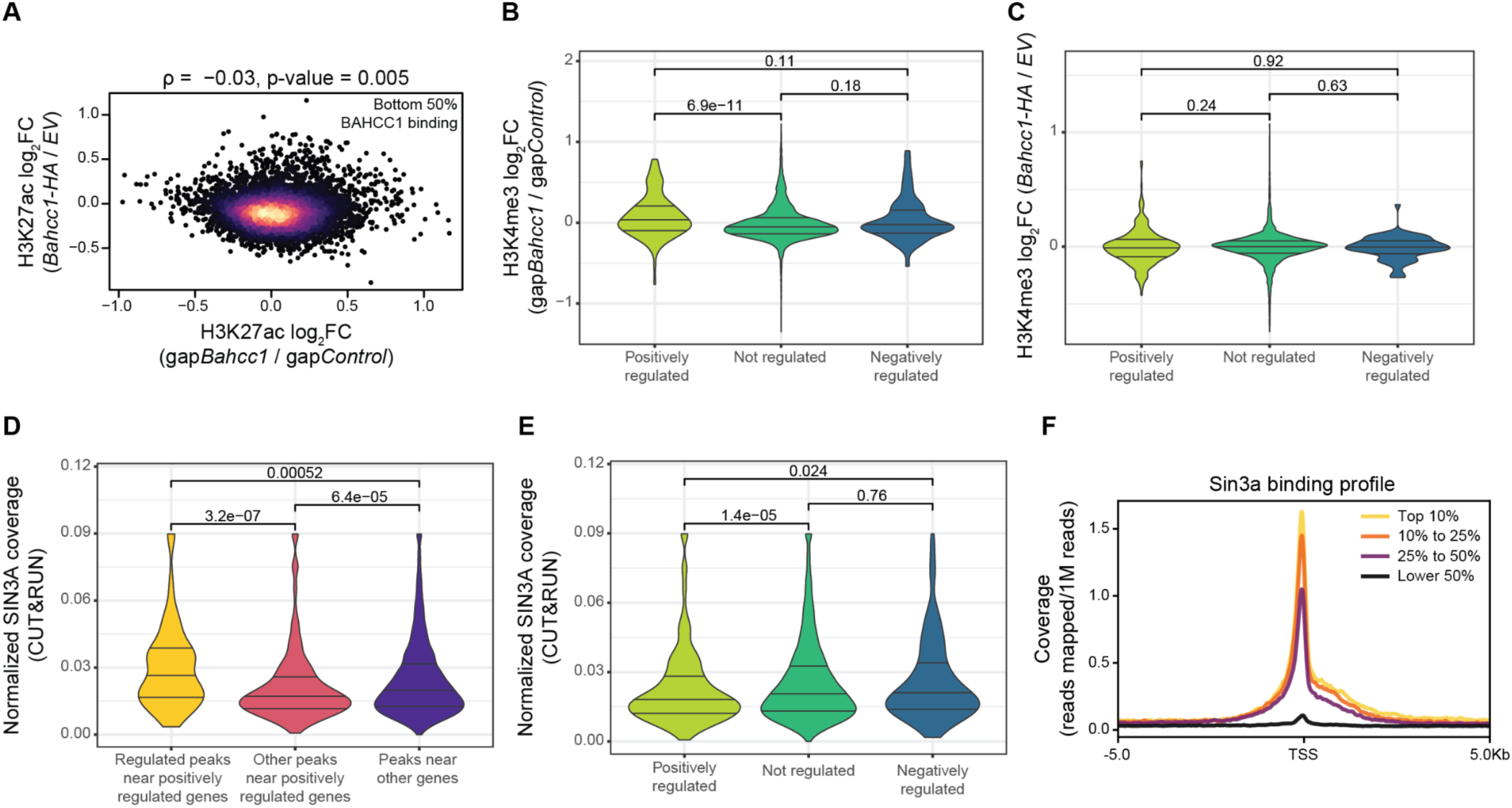
BAHCC1 positively affects gene expression. **(A)** Correspondence between changes in H3K27ac after *Bahcc1* KD and OE for the peaks with the lower 50% average coverage of BAHCC1 CUT&RUN reads. Color indicates local point density. **(B)** Changes in H3K4me3 at peaks near the BAHCC1-regulated genes in *Bahcc1* KD cells. **(C)** Same as in (D), but after *Bahcc1* overexpression. **(D)** Normalized SIN3A CUT&RUN signal over the H3K27ac regulated peaks associated with the positively regulated genes. **(E)** Normalized SIN3A CUT&RUN signal over H3K27ac peaks near positively, negatively, and not regulated genes. **(F)** Metagene profile around TSS of the normalized SIN3A CUT&RUN signal, stratified by gene expression levels in N2a cells. In all violin plots, the three lines represent the median, first and third quartiles, and significance of the different comparisons was calculated by using a Mann-Whitney test.

**Supplementary Figure 5.**
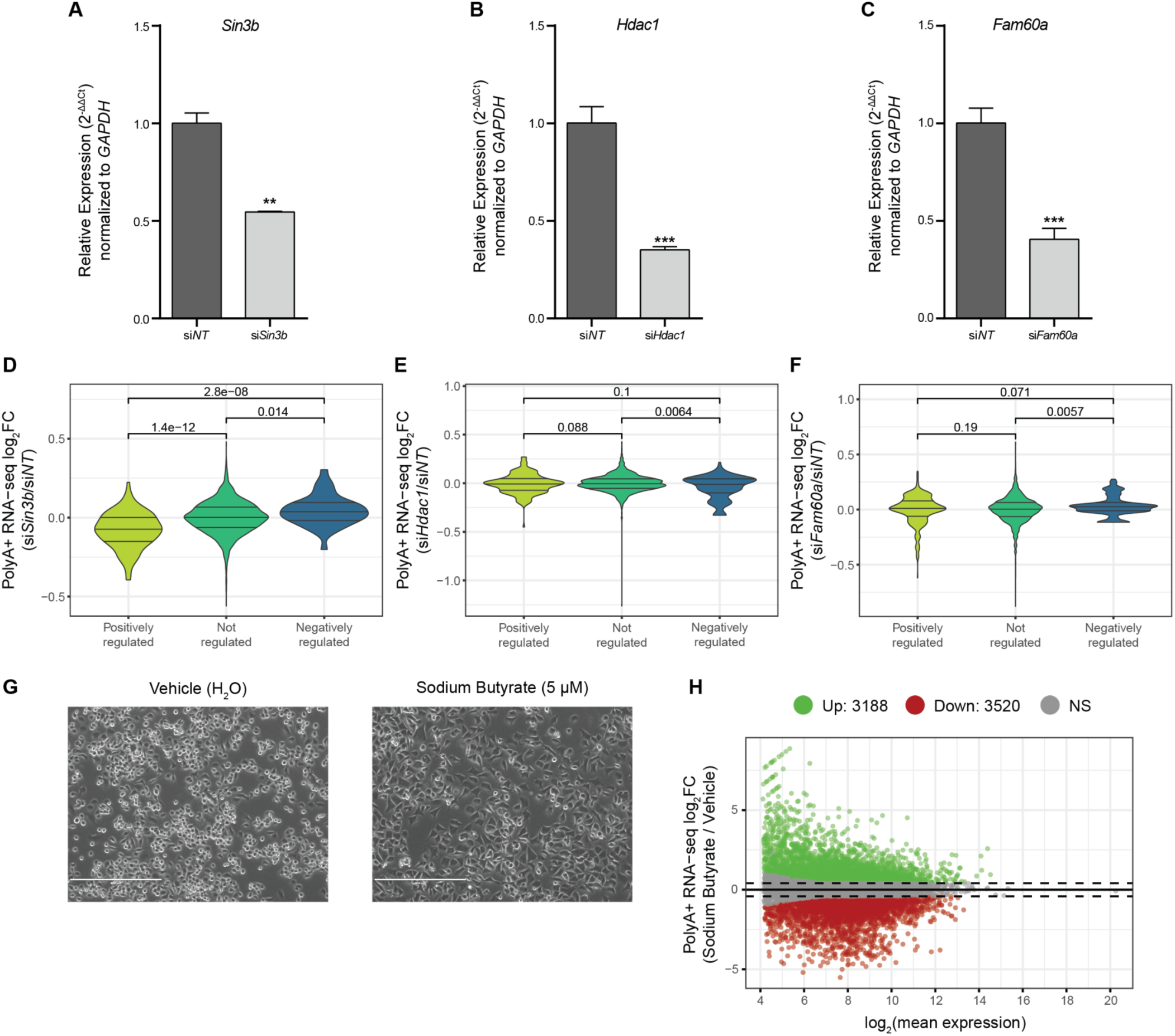
BAHCC1 and different SIN3-HDAC subunits have opposite effects on gene expression. (A-C) RT-qPCR upon depletion of *Sin3b* (A), *Hdac1* (B), and *Fam60a* (C) with siRNAs to assess KD efficiency. **(D-F)** Changes in gene expression upon *Sin3b* (D), *Hdac1* (E), and *Fam60a* (F) KD for the genes positively and negatively regulated by BAHCC1. **(G)** Representative brightfield images of N2a cells after 24 hr of vehicle (left) or sodium butyrate (right) treatment (10X magnification). **(H)** MA plot showing the genome-wide gene expression changes after 24hrs of treatment with sodium butyrate. All experiments were performed in n = 3 biological replicates, with the error bars in the barplots representing the standard deviation, P > .05 = ns; < 0.05 = *, < .01 = **; < .001 = *** (two-tailed Student’s t-test). In all violin plots, the three lines represent the median, first and third quartiles, and significance of the different comparisons was calculated by using a Mann-Whitney test. Genes were considered to be differentially expressed when adj. P-value < 0.05 and |log_2_FC| > 0.41 (corresponding to a change of 33%).

**Supplementary Figure 6.**
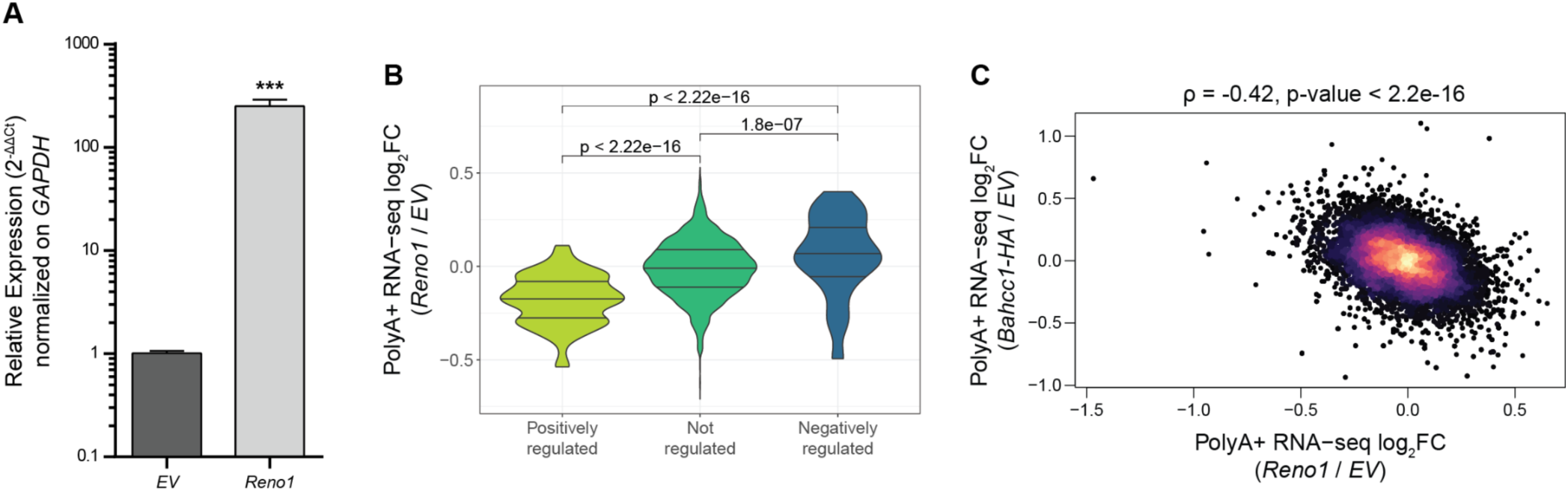
*Reno1* overexpression has opposite effects than *Bahcc1* upregulation. **(A)** RT-qPCR upon transient transfection of a plasmid encoding *Reno1* cDNA to assess OE efficiency. **(B)** Changes in gene expression upon *Reno1* KD with GapmeRs for the genes positively and negatively regulated by BAHCC1. **(C)** Correspondence between changes in gene expression after *Bahcc1* and *Reno1* OE in (-)RA N2a cells. Color indicates local point density. All experiments were performed in n = 3 biological replicates, with the error bars in the barplots representing the standard deviation, P > .05 = ns; < 0.05 = *, < .01 = **; < .001 = *** (two-tailed Student’s t-test). In all violin plots, the three lines represent the median, first and third quartiles, and significance of the different comparisons was calculated by using a Mann-Whitney test.

